# Biofilm Vertical Growth Dynamics Are Captured by An Active Fluid Framework

**DOI:** 10.1101/2025.03.17.643762

**Authors:** Raymond Copeland, Peter J. Yunker

**Affiliations:** Georgia Institute of Technology

## Abstract

Bacterial biofilms, surface-attached microbial communities, grow horizontally across surfaces and vertically above them. Although a simple heuristic model for vertical growth was experimentally shown to accurately describe the behavior of diverse microbial species, the biophysical implications and theoretical basis for this empirical model were unclear. Here, we demonstrate that this heuristic model emerges naturally from fundamental principles of active fluid dynamics. By analytically deriving exact solutions for an active fluid model of vertical biofilm growth, we show that the governing equations reduce to the same form as the empirical model in both early- and late-stage growth regimes. Our analysis reveals that cell death and decay rates likely play key roles in determining the characteristic parameters of vertical growth. The active fluid model produces a single, simple equation governing growth at heights above and below the diffusion limit; surprisingly, this “full” expression is simpler than the heuristic. With this theoretical basis, we explain why the vertical growth rate reaches a maximum at a height greater than the previously identified characteristic length scale. These results provide a theoretical foundation for a simple mathematical model of vertical growth, enabling deeper understanding of how biological and biophysical factors interact during biofilm development.

## 1 Introduction

Biofilms are surface-attached, crowded communities of microbes that can grow in two directions: horizontally across the surface or vertically above the surface[1, 2, 3]. While horizontal growth and range expansion has been more extensively studied ([4, 5, 6, 7]), a growing body of evidence suggests that vertical growth is a crucial aspect of bacterial behavior ([8, 9, 10, 11, 12]). It has even been shown that vertical growth can impact horizontal growth, implying that the expansion of a colony can only be fully understood by considering both phenomena ([13]). Additionally, in some environments, confinement can prevent horizontal growth while allowing vertical growth. Therefore, understanding biofilm growth requires a comprehensive examination of vertical growth dynamics.

Our understanding of vertical growth has been limited in part by its complexity. Nutrients diffuse into the colony through its interfaces, and then are uptaken by microbes. Thus, in a single vertical direction, bacteria experience many different microenvironments [14, 15]. There are many approaches to modeling such systems, cellular automata-based models [16, 17, 18], individual based models [4, 19, 20, 21], or to treat the microbes as an active fluid. Models can even mix aspects of these [22, 23]. In active fluid models, the microbes are essentially growing and energy-consuming particles, characterized by viscous interactions, which allows for the analytical modeling of biofilms based on fluid principles [24, 25]. Prior research has modeled the expansion of biofilms via nutrient diffusion, uptake, and growth as an active fluid [26, 27, 28, 29, 30, 31, 32].

A recent paper, Bravo et al. [15], proposed and experimentally validated a simple heuristic model which we will refer to as the interface model:

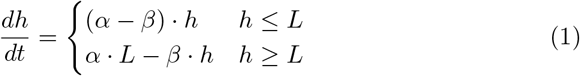

While this model captured the empirical behavior of nine different species, including prokaryotes and eukaryotes, gram negative and gram positive bacteria, and cells with a wide range of shapes and sizes, it was proposed based on empirical data, and justified term-by-term, rather than derived from fundamental dynamics. Thus, the underlying biophysics behind vertical growth remains unclear.

Further, the interface model was unable to explain multiple empirical observations. In particular, the vertical growth rate consistently reached a maximum at biofilm heights greater than the characteristic growing region of length *L*. This effect has also been observed at the edge of growing colonies [13]. However, without a pure theoretical foundation on which to develop hypotheses, it is challenging to investigate the deviations between the model and empirical observations.

Here, we show that the interface model from Bravo, et al.[15], arises directly from a commonly used active fluid model of biofilm growth. We derive the interface model from exact analytical expressions of the active fluid model, and then we fit the experimental data from Bravo, et al. to both the active fluid model and the interface model, showing that the heuristic model is nearly as good. Moreover, we identify that the discrepancies between the interface model and the observed maximum growth rates are effectively captured by the active-fluid model. Thus, we conclude that vertical growth can be accurately modeled using the simpler heuristic framework proposed by Bravo et al. [15], with minimal deviation.

## 2 Active fluid model

Here we develop a one-dimensional model for vertical biofilm growth, with the goal of producing a model in alignment with experimental observations. We initiate this process by deriving a model for growing “bio-fluids” based on the principles of active fluid dynamics. The derivation here closely follows the methodologies established in previous works [30, 27, 28, 22]. We start with the mass continuity equation, which accounts for the flow of various mass, *m*, of species *i* = 1, 2, …*n*:

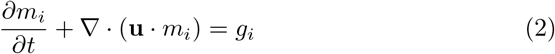

where **u** is the velocity field of the masses and *g*_*i*_ is the growth function for mass species *i*; these can be functions of the other masses or of substrates (as we will later see). As we are currently interested in the growth of a single strain colony, the “bio-fluids” in this model are simply living bacteria; they make-up the domain of the biofilm. As they can be treated as incompressible (they are mostly made of water[33]), we can re-frame the equations away from mass densities, instead replacing them with volume densities *N*_*i*_ = *m*_*i*_*/ρ*; we also replace the growth functions with volume growth functions 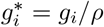:

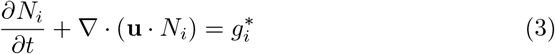

Note this approach also allows for the following constraint:

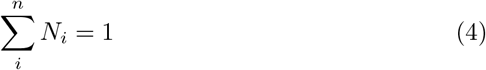

It is next assumed that each of these bio-fluids can be treated as homogeneous, viscous, and incompressible with constant density flowing through a porous environment, satisfying Darcy’s law; this allows us to model the global velocity field with a scalar potential, i.e., the pressure field *p* in (5):

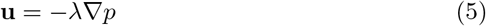

We can now take the volume density continuity equation (3) and sum across bio-fluids, apply the constraint equation 4, expand the momentum divergence (5), and make use of the derivative and sum being linear operators:

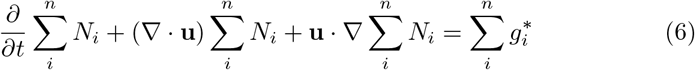

Combining equation 4, equation 5, and equation 6 we are left with an equation which with appropriate choice of boundary conditions allows us to propagate the system:

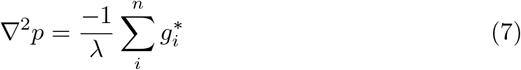

Rewriting equation 3 with equations 7 and 5 leads to a broad expression for a bio-fluid’s propagation:

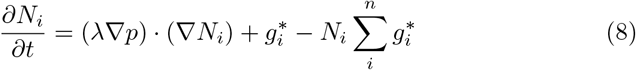

This framework, as detailed in previous studies [30, 28], is well-established. However, to adapt it to our specific system, it is necessary to define the growth functions for each of our bio-fluids. Notably, these terms indicate the increase or decrease of the bio-fluid volume; hence, a positive term signifies volume creation, while a negative term indicates volume loss. We begin by considering a single bio-fluid (*N*_1_) which grows in proportion to both its concentration and the availability of a diffusing resource at a specific location (*R*, see equation 9); it also experiences decay at rate *β* (equation 10). The resources diffuse and are up-taken by the living biomass. The decay rate *β* signifies the rate at which the bio-fluid volume decreases, not the disappearance of the mass itself, but rather the reduction in volume occupied by the bio-mass. This decay rate comes from cell death and the subsequent “quick” breakdown of dead cell structure, discussed below. T

It is well established that bacteria exhibit characteristic death rates, and it is crucial to incorporate such dynamics in our models [34, 35]. Additionally, under conditions of nutrient scarcity (far from diffusing nutrients), bacteria have been observed to engage in programmed cell death [36]. In our model, the rate *β* is the the rate at which microbes die, and we assume it is substantially slower than the rate at which the residual volume of these cells is washed away or broken-down. This assumption simplifies the model by reducing the number of parameters, without significantly altering the dynamics under reasonable expectations.

To illustrate that this is a reasonable assumption, consider another bio-fluid, (*N*_2_), consisting of dead cells that are structurally intact, thus their volume still impacts the living bio-fluid *N*_1_. This material’s volume would increase at a rate *βN*_1_ (due to death of living cells) and decrease at some rate *µ*, so 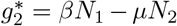. The rate *µ* encompasses all processes that remove the volume of non-growing material, including the diffusion of water originally within the cells out of the biofilm and the breakdown of organic material by other cells [37]. Given that the majority of cell volume is water, tracking the water from the dead bio-mass becomes the critical component to consider. Thus, similar to how nutrient diffusion at rates much higher than biofilm growth, the ‘breakdown’ rate *µ* would be significantly higher than *β*. Therefore, under this fast ‘breakdown’ approximation, both the growth of 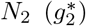 and its relative value would be near zero, so 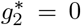 and *N*_2_ ≪ *N*_1_, and from the first expression we arrive at *βN*_1_ ≈ *µN*_2_. From this, we see that the system dynamics would remain basically unchanged, as the sum of the growth functions governs the movement of the bio-fluid, equation 7, and if *βN*_1_ ≈ *µN*_2_ and *N*_2_ ≪ *N*_1_, the sum of the growth functions would mirror that of our current model.

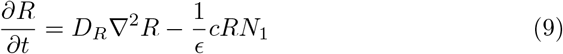

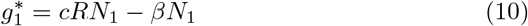

Additionally, while alternative common growth functional forms related to resource consumption, such as the Monod equation, could certainly be utilized, the form we have adopted facilitates the solution of the system. The use of an unbounded maximum growth rate as *R* increases could potentially introduce complications; however, the Dirichlet boundary condition of our resources at *R*^*^ stabilizes *R* to be at most *R*^*^ (so long as the colony does not drop suddenly in height). Typically, growth is defined explicitly by a predetermined maximum growth rate *α*, and we can still impose such a maximum by aligning it with a realistic values through the use of constant *c* in (9), such that *α* = *cR*^*^. It is important to note that *ϵ* is a unit-less constant that ensures mass conservation in our model, derived from [38].

As our analysis is confined to a single dimension (*z*), all gradients and spatial derivative operators are exclusively with respect to *z*. The challenge arises with the boundary conditions. As the colony dynamically grows or contracts, the top boundary shifts, complicating the definition of boundary conditions and rendering them computationally infeasible to implement. However, we can define the height over time as it is solely a function of the velocity field **u**:

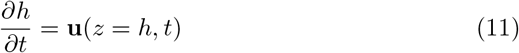

We now define our boundary conditions as: 1) the gradient of pressure must be zero at *z* = 0, and 2) a Dirichlet boundary (*p* = 0) at the top of the colony (*z* = *h*) in line with previous work [24]. The resource *R* boundary conditions are a source term at *z* = 0 such that its concentration is constant, and the gradient of *R* is zero at the top of the colony. Thus, resources flow freely from the bottom of the colony, and they reflect off the top of colony back into it:

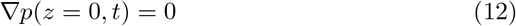

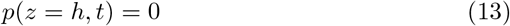

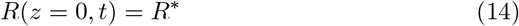

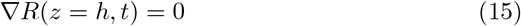

## 3 Derivation of heuristic model

We now derive a solution for the height of the colony over time from the active fluid model from section 2. There is one bio-fluid (*N*_1_), which is defined by the growth function in equation 10. So, to find the height over time, we must know the velocity field at the top of the colony, which means we must know the pressure gradient, which means we must know the solution of the growth functions, which means we must know the solution for how *R* and *N*_1_ are distributed in space. Given the fast diffusion of resources, we take the quasi-static approximation such that *∂R/∂t* = 0. Thus, the resource curve takes on a classic exponential form:

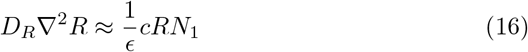

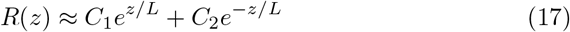

Using the boundary conditions defined in equations 14 and 15, we arrive at the following expressions for our unknown constants. Note that we took *N*_1_ to be = 1 and constant across space; this is a valid result given that there is only one bio-fluid under the constraint of equation 4. We find a characteristic length scale *L* (equation 18) at which resources decay; it is proportional to the diffusion of resources and inversely to uptake rate. This functional form of this length scale is similar to those in some horizontal models [39].

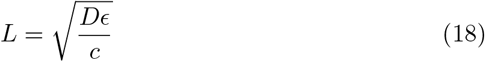

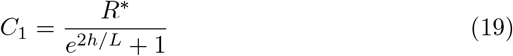

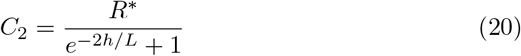

Thus, as *h* grows, the contribution of the positive exponential term decays ; however it is significant in early times when *h* is low, see Fig2B. Physically, the positive exponential term represents the accumulation of resources at the top boundary of the biofilm, and their subsequent diffusion back into the biofilm bulk. This is illustrated in Fig2A.

At this point it is convenient to undergo a coordinate transformation according to equation 21. This does not change any result we already have derived so far, but it will change our derivative functions going on according to the equations below ( 21,22, 23, 24, and 25). This transformation allows us to rigorously define the domain *z*^′^ ∈ (0, 1) and our boundaries can be applied computationally. Similar coordinate transformations were used in [27, 28].

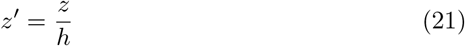

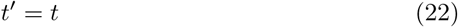

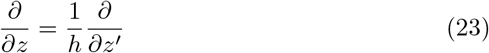

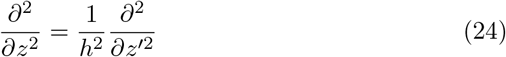

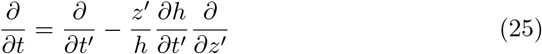

With this coordinate transformation we can express the growth of our bio-fluid on a defined domain, see equation 26. Note, we replaced *cR*^*^ with *α*, and we are now in *z*^′^ coordinates. It will become clear later that this *L* is the same *L* as in the interface model.

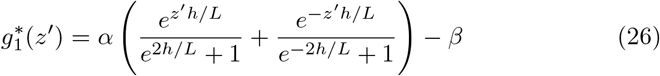

Now we can combine our expression for the growth of the bio-fluid with our expression for pressure (equations 7 and 10) which yields the following expression for the Laplacian of pressure (see equation 27). The extra factor of *h*^2^ appears from the coordinate transformation.

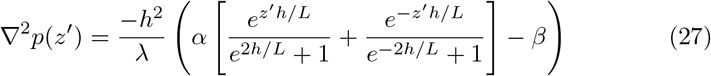

As the Laplacian is in not a function of itself and only spatially defined by *z*^′^, we can integrate across the whole domain, and then use the boundary conditions to find the velocity field at the top of the colony (see equation 28).

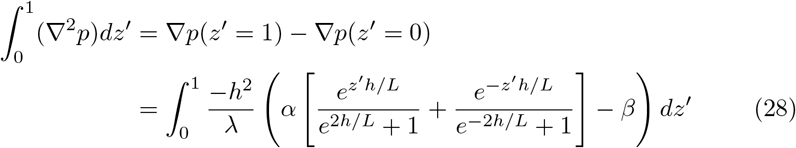

We integrate this, use our boundary condition stating that the gradient of pressure must be zero at *z*^′^ = 0, and then plug it into equation 11 in the modified coordinates (which cancels out another factor of h) yielding:

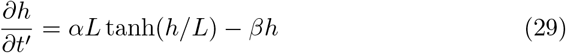

Thus we recover the interface model described in Bravo et al. The active growth regime of length *L* is the characteristic length of diffusing resources as suggested in the study. At *h* ≫ *L*, we are left with:

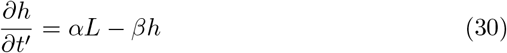

and at *h* ≪ *L*:

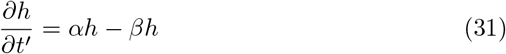

Using the expression in 29 allows us to find the height at which the growth rate is maximized by taking the derivative with respect to *h* and setting the expression to zero:

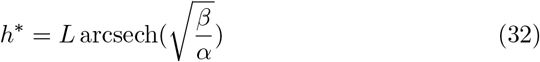

## 4 Comparison between experiments and exact and approximate numerical predictions

We now compare the interface model (as outlined in Bravo, et al., [15]) and the full analytical expression (equation 29), referred to as the fluid model, to experimental data (originally published in [15]). Figure 1A illustrates the best fits of both the interface and fluid models against the height-over-time curve of an A. veronii colony, revealing that the two models are virtually indistinguishable. Furthermore, Figure 1B highlights the similarity in form between the fluid and interface models, though the rationale behind the fluid model’s form is more transparent, and it improves upon the interface model’s insights. To comprehensively demonstrate that the fluid model is as robust as the interface model, we fitted the height-over-time curves of all nine strains outlined in [15], as presented in Fig. 3. As observed previously, the fits between the two models are indistinguishable. This similarity is expected, given that both models are grounded in the same fundamental forms and share the same number of parameters. We also find that the fit parameters are all very similar to each other as seen in Fig 1A.

**Figure 1:**
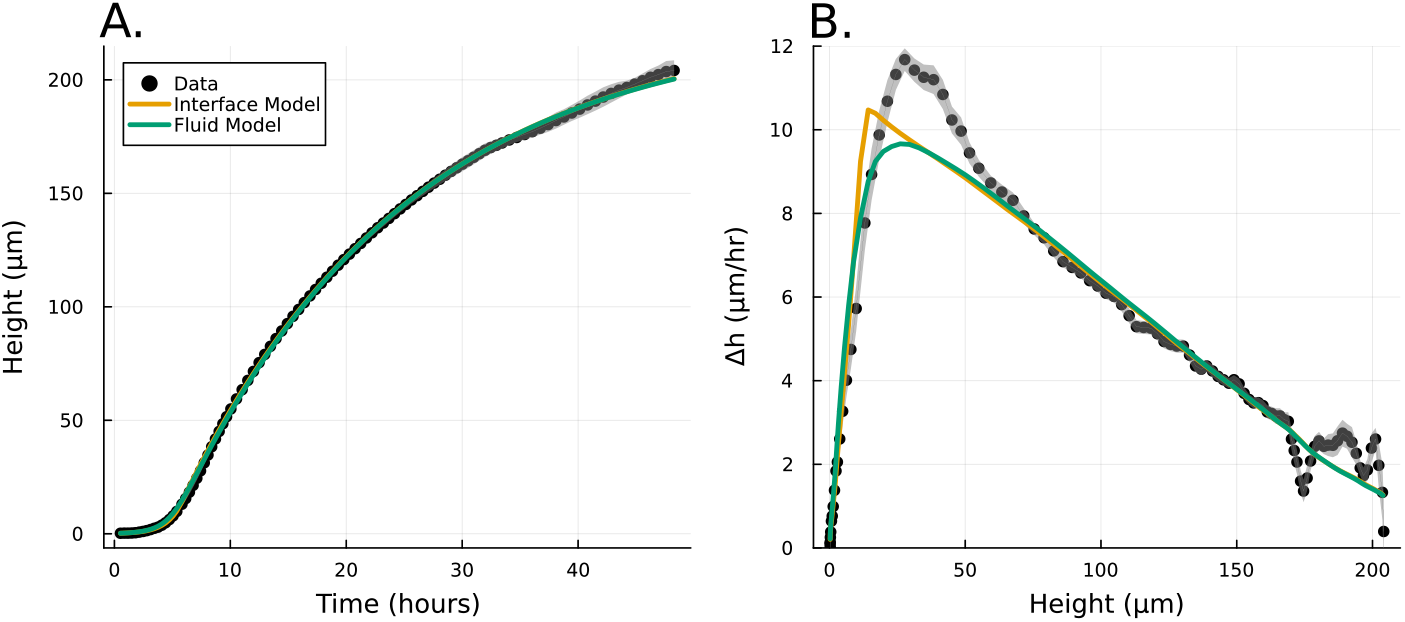
A. Experimental data from Bravo et al. [15] depicting the growth of A. veronii colony height over time, measured using an interferometer. The interface model was applied using equation (1) and the fluid model using equation (29). Fit parameters for the fluid model are *α* = 0.86 hr^−1^, *β* = 0.051 hr^−1^, and *L* = 13.17 *µm*; for the interface model, *α* = 0.82 hr^−1^, *β* = 0.050 hr^−1^, and *L* = 13.86 *µm*. B. This panel shows the slope of the height over time curve as a function of height, with experimental data again sourced from Bravo et al. [15]. The fluid model is represented by the green line and the interface model by the orange line, as in panel A.

**Figure 2:**
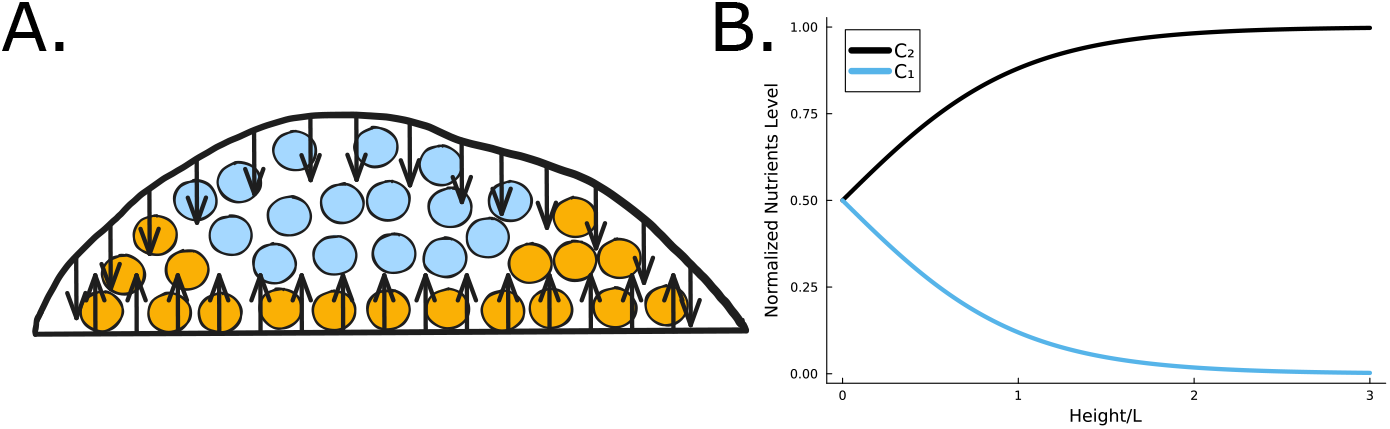
A. When the total height of the colony is small, the reflective top surface of the biofilm allows for more growth (orange cells) than when the colony is large, and the cells beyond *L* can no longer grow (blue cells). B. *C*_1_ is the contribution of the exponentially growing term in equation 17 while *C*_2_ is the exponentially decaying term.

**Figure 3:**
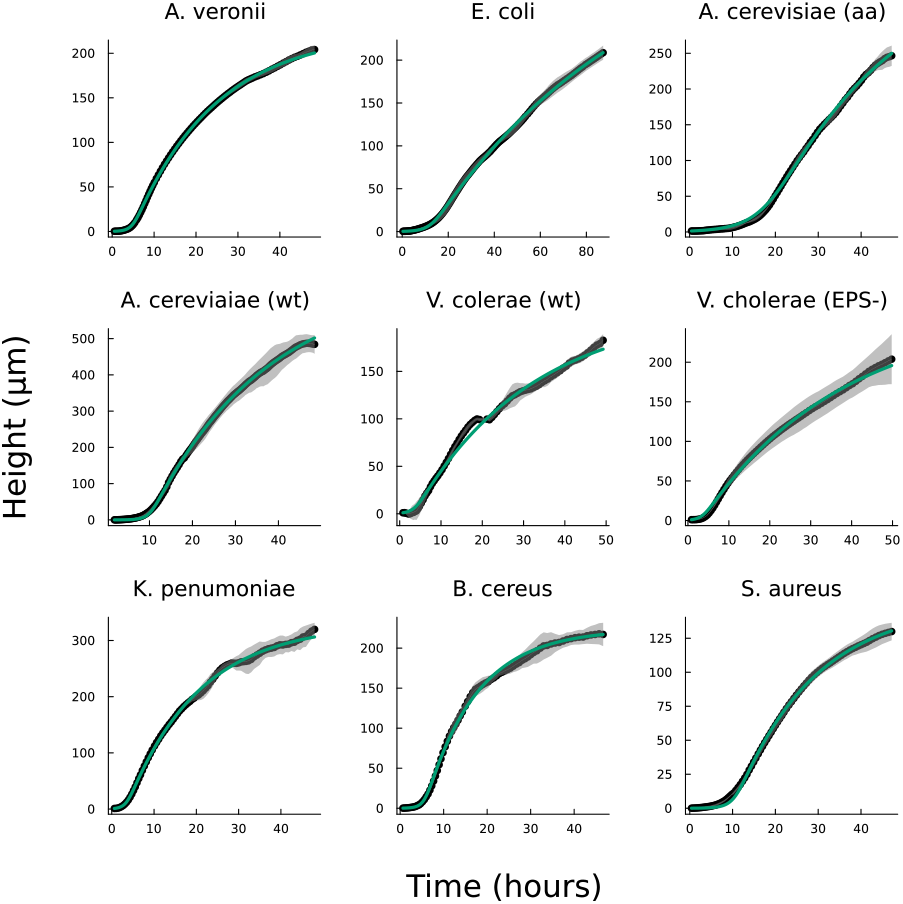
Displayed is the fluid model fit to the same experimental data from Bravo et al. [15] using equation (29). The fit is as precise as the interface model detailed in (1) with similar differences in fit parameters as shown in Fig. 1.

## 5 Resolving Discrepancies in Models of Biofilm Growth Dynamics

Note, our objective is not to present a model that better fits the experimental data, but rather to validate that the broad functional form adopted is appropriate and captures a previously unaddressed observation. In particular, the interface model proposes a characteristic height, *L*, which marks the transition from the actively growing regime to the nutrient-depleted regime, a transition from mostly growth to mostly decay. This would suggest that in every cellular layer above height *L*, the rate of decay exceeds the rate of growth. Thus, height *L* should represent the level at which biofilm growth reaches its maximum growth. However, experiments consistently show that the growth rate peaks (denoted *h*^*^) at a biofilm height *h*^*^ *> L*; specifically, as illustrated in Fig 1B and Fig 4A, *L* does not align with *h*^*^.

**Figure 4:**
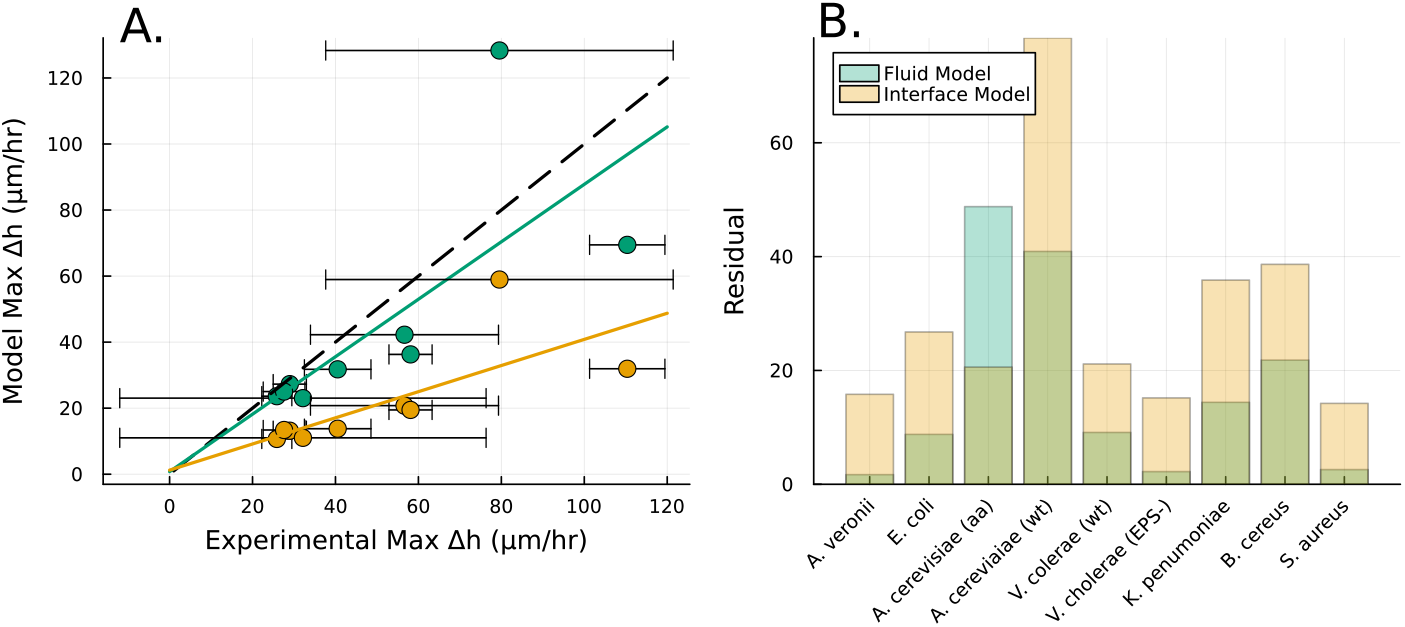
A. The height *h*^*^ at which Δ*h/*Δ*t* is maximized, as calculated from equation (32), is plotted in green against the experimental peak in Δ*h/*Δ*t* versus *h* from all data shown in Fig. 3. In orange, the *L* fit from the interface model is plotted against the same experimental peaks. B. The residuals, representing the distance from the experimental peak in A., are displayed for each strain to illustrate how the fluid model more accurately captures this phenomena in all but one strain.

The fluid model provides a framework to resolve this discrepancy. The best fit *L* values in both models are notably similar, which can be understood through the work presented in the previous section. While *L* represents the characteristic length at which resources are depleted in both models, equation 30 shows that *L* only represents the size of the growing region when the colony achieves sufficient height (*h* ≫ *L*); if *h* ∼ *L*, then the thickness of the active growing region is greater than *L*. This phenomenon arises from the boundary condition on the upper surface of the colony where nutrients are reflected back inside the biofilm; this prevents a simple characteristic decay of nutrients, as demonstrated by equations (19) and (20). The nutrients are reflected back, so at low *h* the cells have an abundance of nutrients (see Fig. 5A and Fig. 2B). Thus, as the biofilm grows, the reflective surface becomes farther from the nutrient source (with increasing *h*), the taller biofilm develops a distinct characteristic depth *L* of metabolically active cells. This phenomenon is qualitatively illustrated in Fig. 2A. Fig 5A demonstrates that when the colony initially attains height *L*, the nutrient concentration at *z* = *L* can be approximately 70% higher due to the boundary effect, compared to later where *h* exceeds *L*, and the two converge. This interaction results in a more gradual transition in our final expression, and it facilitates the theoretical determination of *h*^*^ through equation (32). Experimentally measured *h*\* values are much more aligned with the fluid model predictions than with the interface model *L* (Fig. 4A and B).

**Figure 5:**
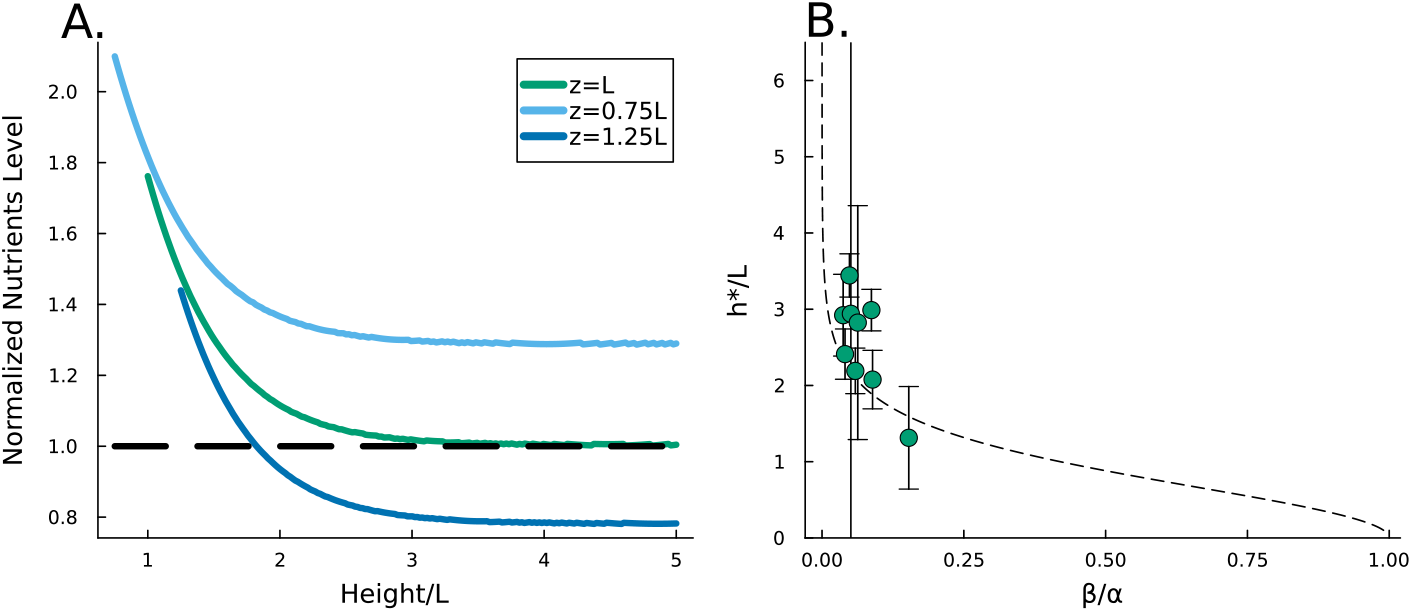
A. Shown is the available nutrient concentration at three different *z* distances as the height of the colony grows; we see that as the colony grows the nutrient concentration decreases. The black dashed line represents the steady state concentration at *z* = *L*, so we see that at low *h* the growing region is greater than *L*. B. The ratio *h*^*^*/L* , where *h*^*^ is the experimental height at which Δ*h/*Δ*t* is maximized and *L* is the fit parameter from equation (29), is plotted against the fit parameters *β/α* as green circles. The dashed line represents equation (32). We observe that the strains from Bravo et al. [15] occupy a closely aligned regime within the parameter space, and the fluid model closely approximates the predicted *h*^*^*/L*.

An alternative explanation could be reached from figure 4A which demonstrates a nearly linear positive correlation between *L* and experimental *h*^*^, this relationship might suggest there is residual growth beyond the characteristic nutrient decay length *L*, consistent with the persistent nature of exponential decay curves. In essence, *L* and *h*^*^ may only be a factor apart, akin to one more, albeit smaller, standard deviation of nutrients. To explore this, we turn to the dimensionless ratio *h*^*^*/L*, which represents the factor by which a colony’s maximum growth exceeds its characteristic length *L*. The fluid model predicts this ratio via equation (32); it shows good agreement with experimental data, with seven of nine strains falling within one standard deviation of the fluid model’s prediction (Fig 5C). Furthermore, this analysis reveals that the examined strains occupy a narrow region in parameter space (specifically, *β/α*). Thus, while the apparent proportionality between *h*^*^ and *L* might suggest a universal relationship, this correlation may only hold when the ratio *β/α* remains relatively consistent across strains.

## 6 Discussion

This study presents a theoretical foundation for an empirically established model of vertical biofilm growth. Through rigorous mathematical analysis, we demonstrate that the heuristic “interface model” previously developed in [15] emerges naturally from fundamental principles of active fluid dynamics.

Our analysis suggests that the *β* parameter in the heuristic model could primarily correspond to cell death with fast ‘breakdown’ of the dead cell structure. While additional factors—including viscous relaxation and extracellular matrix degradation—likely contribute to this term, we demonstrated here that cell death and lysis alone, under fast ‘breakdown’ conditions, can account for the observed *β* behavior. This finding elucidates why the relatively parsimonious heuristic model exhibits robustness across diverse microbial species.

We also validated our fluid model against experimental data from the preceding study. The Fluid model demonstrates exceptional agreement in height-versus-time across nine distinct microbial strains. Furthermore, the model provides a theoretical framework explaining why growth rates achieve a maximum at heights exceeding the characteristic length *L*—a phenomenon that remained unexplained by the previous heuristic model. Our analysis reveals that boundary conditions at the colony’s upper surface enhance growth potential in shorter colonies. This insight suggests the possibility of analogous effects in horizontal growth at colony edges, where the cells are always exponentially growing and have two reflective bounds [13].

Intriguingly, the analysis presented here suggests that a relatively small range of *h*^*^*/L* because the experiments happened to have a narrow range of *β/α*. Future research could explore if this narrow range of *β/α* was due to random chance, or if most microbes grown in the lab fall in this range. However, it is important to note that quantifying cell death rates within dense biofilm structures presents substantial experimental challenges.

An alternative approach to validating the active fluid model would involve extending the nutrient dynamics to two or three spatial dimensions as previous models typically assume logistic growth vertically. Concordance between such an extended model and experimental observations would provide compelling evidence for the model’s fundamental robustness, rather than mere consistency with existing data.

